# A single mutation attenuates both the transcription termination and RNA-dependent RNA polymerase activity of T7 RNA polymerase

**DOI:** 10.1101/2021.01.11.426313

**Authors:** Hui Wu, Ting Wei, Rui Cheng, Fengtao Huang, Xuelin Lu, Yan Yan, Bingbing Yu, Xionglue Wang, Chenli Liu, Bin Zhu

## Abstract

Transcription termination is one of the least understood processes of gene expression. As the prototype model for transcription studies, the single-subunit T7 RNA polymerase (RNAP) was known to response to two types of termination signals, while the mechanism underlying such termination especially the specific elements of the polymerase involved in is still unclear, due to the lack of a termination complex structure. Here we applied phage-assisted continuous evolution to obtain variants of T7 RNAP that can bypass the typical class I T7 terminator with stem-loop structure. Through *in vivo* selection and *in vitro* characterization, we discovered a single mutation S43Y that significantly decreased the termination efficiency of T7 RNAP at all transcription terminators tested. Coincidently, the S43Y mutation almost eliminates the RNA-dependent RNA polymerase (RdRp) of T7 RNAP without affecting the major DNA-dependent RNA polymerase (DdRp) activity of the enzyme, indicating the relationship between transcription termination and RdRp activity, and suggesting a model in which the stem-loop terminator induces the RdRp activity which competes with the ongoing DdRp activity to cause transcription termination. The T7 RNAP S43Y mutant as an enzymatic reagent for *in vitro* transcription reduces the undesired termination in run-off RNA synthesis and produces RNA with higher terminal homogeneity.

## INTRODUCTION

Transcription termination of RNA polymerase (RNAP) is a crucial step for regulation of gene expression by triggering the release of RNA transcripts and RNAPs from the transcription complex. RNA polymerases in most organisms have complex subunit compositions, structure and regulatory factors (1–3). And the transcription termination is a highly intricate and dynamic process, making related biochemical and structural studies extremely difficult. Among stages in transcription (initiation, elongation and termination), the molecular mechanism of transcription termination remains the least understood (4,5). Even for the single-subunit RNAP from bacteriophage T7, of which the initiation and elongation complex structures had been solved (6,7) and greatly improved the understanding of transcription mechanism, a snapshot of its termination is still missing, as no termination complex structure available.

Two types of termination signals were known to induce transcription termination of T7 RNAP. The class I terminators encode nascent RNA with a stable stem-loop structure followed by a run of U residues, such as T7 TΦ found at the late region of T7 genome (8–10). This type of terminator is analogous to the rho-independent terminators of bacterial RNAPs. Indeed, T7 RNAP was reported to terminate at several of such bacterial terminators (11,12). The second type signals encode RNA with a common sequence (5’-AUCUGUU-3’) and T7 RNAP terminates at 6-8 nt downstream of this sequence. The class II terminators are strictly sequence-specific as any base change would abolish the termination (13,14). Class II terminator was first identified in a cloned human preproparathyroid hormone (PTH) gene. Additional class II signals were also found in a cDNA copy of vesicular stomatitis virus (VSV), adenovirus DNA (Adeno5), and near the right end of the concatemer junction (CJ) of replicating T7 DNA (9,13). The mechanism underlying the class-II-signal induced termination is especially elusive since these short terminator sequences do not indicate any distinct secondary structure. To date, no direct information has been obtained concerning structure changes during the termination process of T7 RNAP.

In addition to being a research model for transcription mechanism, T7 RNAP also serves as a popular enzymatic tool for *in vitro* transcription (IVT) (15–17). IVT for RNA synthesis is widely used in RNA related research and applications especially the recently booming RNA-based therapeutics such as siRNA and mRNA drugs and vaccines (18–20). However, various kinds of byproducts generated by T7 RNAP during IVT, including short RNAs produced by abortive initiation (21), undesired terminated transcripts caused by termination signals (10), 3’ terminal extended products resulted from the RNA-dependent RNA polymerase (RdRp) activity (22,23) may activate the vertebrate innate immune system to impede RNA-based therapeutics (24). Although methods such as high performance liquid chromatography (HPLC) are available to remove certain RNA byproducts from IVT (25), they are usually not optimal for large scale RNA purification and will increase production complexity and cost. It is advantageous to develop enzymatic reagents to reduce the undesired RNA byproducts from IVT, and to improve the rapidly developing RNA-based applications and therapeutics.

In this study, we set up a phage-assisted evolution system (PACE) to select T7 RNAP mutants that can bypass the typical stem-loop T7 terminator TΦ. We aimed to reveal the elements in the RNAP crucial for transcription termination and improve the understanding of the mechanism underlying the termination process. As a result, we found that a single serine to tyrosine mutation (S43Y) significantly decreased the termination efficiency of all terminators tested without affecting the major DNA-dependent RNA polymerase (DdRp) activity of the enzyme. Unexpectedly, the S43Y mutation almost abolishes the RdRp activity of T7 RNAP, resulted in higher 3’ homogeneity of IVT products. Based on these results, a model was proposed with regarded to the relationship between the termination at class I terminator and the RdRp activity of T7 RNAP. This study provided insights into the mechanism by which the single-subunit RNAP responds to termination signal in nascent RNA.

## MATERIALS AND METHODS

### Materials

Oligonucleotides were from Genecreate. DNA purification kits were from Axygen. RNase Inhibitor and DNA marker were from Thermo Scientific. Low Range ssRNA Ladder and RNA purification kits were from New England Biolabs. HPLC reversed-Phase column PLRP-S was from Agilent. PrimeSTAR Max DNA Polymerase was from TaKaRa. Ni-NTA resin was from Qiagen. Competent *E. coli* DH5α and *E. coli* BL21 were from TransGen Biotech. ClonExpress II One Step Cloning kits for vector construction were from Vazyme Biotech.

### Bacterial strains and Plasmids

Plasmids used in this work were amplified in *E. coli* DH5α and enzymes expressed in *E. coli* BL21. The information of bacterial strains and plasmids used for PACE experiments was described in Table S1 (26,27). The plasmids used for IVT templates preparation include pETsnrs-1 (Table S4) carrying T7 promoter and T7 terminator TΦ (5’-CTGCTAACAAAGCCCGAAAGGAAGCTGAGTTGGCTGCTGCCACCGCTGAGCAATAAC**TAGCATAACCCCTTGGGGCCTCTAAACGGGTCTTGAGGGGTTTTTTG**CTGAAAGGAGGAACTATAT-3’, bold sequence forms stem-loop structure, Figure S1), and pUCT7sg-eGFP carrying T7 promoter and eGFP sgRNA coding sequence. Other plasmids with different terminators were constructed by replacing the stem-loop forming region (5’-**TAGCATAACCCCTTGGGGCCTCTAAACGGGTCTTGAGGGGTTTTTTG**-3’) in TΦ with various terminator sequences. The plasmids for IVT templates of various sgRNA were constructed by replacing the first 20 nt of the eGFP sgRNA coding sequence with other guiding sequences (Table S3).

### PACE experiments

*E. coli* S1030 cells carrying mutagenesis plasmid and either of the accessory plasmids (pLATp1 and 2) were used for the PACE experiments. Overnight cultures were 1:100 diluted in fresh LB and shaking cultured until early exponential phase. Then 170 μl of the bacterial culture, 20 μl of the initial selection phages carrying wild-type T7 RNAP gene, and 10 μl of L-arabinose (1 M) were added to each well of a Corning 96-well standard microplate. The rest of the bacterial cultures were temporarily stored at 4°C. Three replicates were performed for either host bacterium. The plate was incubated at 37°C in a Digital Thermostatic Shaker (AOSHENG) with a shaking speed of 1000 rpm. After 3 h incubation, 20 μl of the culture from each well was transferred to a new well, and then 170 μl of the stored bacterial culture and 10 μl L-arabinose were added. The plate was again shaking cultured at 37°C. The culture-and-transfer cycle was repeated, and the culturing time was reduced by 15 minutes every two cycles until it reached 35 min. When the cycles were paused at the end of a working day, samples from the last transfer were collected and centrifuged at 15,000 rpm for 5 min, and the supernatants were collected and stored at 4°C. Bacterial cultures were freshly prepared every day. Supernatant samples containing evolved phages from the last cycle were subjected to double-layer plaque assay and phage clone purification with host cells carrying either of the accessory plasmids (pLATp1, 2). Purified phage clones were used for Sanger sequencing of the T7 RNAP gene and further analyses.

### Phage titer quantification

Phage titer was quantified by double-layer plating plaque assay. Selection phage samples carrying wild-type or mutant RNAP genes were prepared by 10-fold serial dilutions with fresh LB. Overnight cultures of *E. coli* S1030 cells carrying each of the accessory plasmids (pLATp1~3) were 1:100 inoculated into fresh LB and incubated with shaking at 37°C until the OD_600_ was over 0.6. Then 200 μl bacterial suspension and 10 μl phage sample were added to 3.6 ml melt LB containing 0.6 % (w/v) agar, mixed by vortex, and poured into an 8.5-cm Petri dish containing a bottom layer of hard LB agar (1.5 %, w/v). The plates were gently shaken to allow the melt soft agar to spread all over the surface of the bottom agar. After the soft agar layer completely solidified, plates were cultured overnight at 37 °C. Plaques were counted to quantify the titer of the phage samples. Plaque assay was conducted in triplicate. Phage titer obtained using host bacterium carrying pLATp3 was used as the standard for calculating the efficiency of plating (EOP).

### Luminescence reporter assay

*E. coli* S1030 cells carrying each of the reporter plasmids (pLRTp1~3) were used for the luminescence reporter assay. Colonies from freshly streaked agar plates were inoculated into LB broth and cultured overnight. The cultures were then 1:100 inoculated into fresh LB and cultured until mid-exponential phase. OD_600_ was measured with a Thermo Genesys 10S ultraviolet spectrophotometer. Bacteria suspensions were diluted to adjust the OD_600_ to 0.1, and 20 μl of the diluted sample was added into 170 μl LB in each well of a Corning 96-well clear bottom white polystyrene microplate. Selection phage samples carrying wild-type or mutant T7 RNAP genes were prepared by adjusting phage titer to approximately 10^11^ PFU/ml, and 10 μl of the sample was added into each well to make up a total volume of 200 μl. The 96-well plate was then incubated at 37°C in a Digital Thermostatic Shaker (AOSHENG) at a shaking speed of 1000 rpm. After 2 h of incubation, the optical density and luminescence were measured using a Biotek Synergy H1 multi-mode microplate reader. The luminescence of each sample was normalized by OD, and the values obtained using host bacterium carrying pLRTp3 were used as the standard for calculating the relative luminescence. Three independent assays were conducted for the measurement.

### Protein expression and purification

DNA fragments encoding T7 RNAP variants were amplified from the phage genomic DNA and were inserted into pQE82L vector encoding N-terminal His-tag. Vectors pQE82L carrying wild-type (WT) T7 RNAP gene or its variants were transformed into *E. coli* BL21. The bacteria were cultured in 250 mL LB medium containing 100 μg/ml ampicillin at 37°C until they reached an OD_600_ of ~1.2, then the protein expression was induced by the addition of 0.5 mM IPTG and incubation was continued at 30°C for 4 h. The cells were harvested, resuspended in 300 mM NaCl, 20 mM Tris-HCl pH 7.5, 0.5 mM DTT, and lysed by three cycles of freeze-thaw in the presence of 0.5 mg/ml lysozyme. Cleared lysates were separated by centrifugation for 1 h at 18000 rpm, 4°C. His-tagged T7 RNAP mutants were then purified from the lysate using Ni-NTA-agarose column. After equilibrating the column with elution buffer (300 mM NaCl, 20 mM Tris-HCl pH 7.5, 0.5 mM DTT), the RNAP was eluted from the column with 30ml gradient (20-100 mM imidazole) in elution buffer. Then the fractions eluted by different concentration of imidazole were collected and analyzed on SDS-PAGE gels with Coomassie blue staining. The fractions with high purity were then combined and concentrated by Amicon Ultra-15 Centrifugal Filter Units (Millipore). Finally, the concentrated samples were dialyzed three times against a dialysis buffer (100 mM NaCl, 50 mM Tris–HCl, pH 7.5, 1 mM DTT, 0.1 mM EDTA, 50% glycerol, and 0.1% Triton^™^ X-100). Single mutations were introduced to the 43 position of T7 RNAP using ClonExpress II One Step Cloning Kit (Vazyme) and the same purification process as above was applied.

### Transcription assays

To compare the termination efficiency of the T7 RNAP WT and its mutants, we conducted IVT assays (Figure 2B and 2C). Transcription template was amplified through PCR from the plasmid pETsnrs-1 with a pair of universal primers (pETsnrs-F: 5’-ATCAGGCGCCATTCGCCATTCAGG-3’ and pETsnrs-1R: 5’-TCGGTGAGTTTTCTCCTTCATTAC-3’). Reaction mixtures containing 40 mM Tris-HCl pH 8.0, 16 mM MgCl_2_, 2 mM spermidine, 5 mM DTT, 0.2 μM inorganic pyrophosphatase, 1.5 U/μl RNase Inhibitor, 4 mM each of ATP, CTP, GTP, and UTP, 14 nM DNA templates and 50 nM T7 RNAP or its mutants were incubated at 37°C. After incubation for 1 h, DNA templates were removed by DNase I and the transcripts were purified by RNA purification kits (New England Biolabs). Purified RNA was quantified by measurement of OD_260_ using NANOPHOTOMETER (IMPLEN). 400 ng RNA transcripts were mixed with 4 μl 3x denaturing loading buffer (95% formamide, 1 mM EDTA, 0.02% bromophenol blue, 0.02% SDS and 0.01% xylene fluoride) and 7 μl ddH_2_O, heated at 80°C for 2 min, then loaded onto a 1.5% TAE agarose gel. Electrophoresis was performed at 100 V for 45 min and the RNA products were visualized by staining with ethidium bromide.

To further demonstrate the effect of the S43Y mutation of T7 RNAP on termination at terminator TΦ, the reactions were performed at various enzyme concentrations (5.6 nM, 16.7 nM, 50 nM, 150 nM, 450 nM), template concentrations (1.75 nM, 3.5 nM, 7 nM, 14 nM, 28 nM) or reaction time (0.25 h, 0.5 h, 1 h, 2 h). To examine whether the S43Y mutation has a common effect on the class I terminators (Figure 3D), the terminator T7 TΦ was replaced by T3 TΦ, *thr* attenuator, *rrnBT1* terminator or *rrnC* terminator, respectively (Table S2). Transcription templates carrying various terminators were amplified by PCR using a pair of universal primers (pETsnrs-F and pETsnrs-1R), and the IVT reaction was conducted as mentioned above.

To test the effect of T7 RNAP S43Y mutation on class II termination signals, an artificial class II terminator (5’-ATCTGTTTTTTTTT-3’) was inserted into the plasmid pETsnrs-1 by replacing T7 TΦ (Figure 4A). Transcription templates were amplified by PCR using primers pETsnrs-F and pETsnrs-2R (5’-TAAATCAAAAGAATAGACCGAGATAGG-3’), the termination efficiency of the WT T7 RNAP and S43Y mutant at this artificial class II terminator and class I T7 TΦ was compared with the same IVT reaction condition as above. To further verify the effect of S43Y on class II signals, another weaker class II terminator (5’-ATCTGTTT-3’) was introduced into the transcription template of an eGFP sgRNA coding sequence (5’-GGGCACGGGCAGCTTGCCGGGTTTTAGAGCTAGAAATAGCAAGTTAAAATAAGGCTAG TCCGTT**<u>ATCAACTT</u>**GAAAAAGTGGCACCGAGTCGGTGCTTTTTTT-3’ to 5’-GGGCACGGGCAGCTTGCCGGGTTTTAGAGCTAGAAATAGCAAGTTAAAATAAGGCTAG TCCGTT**<u>ATCTGTTT</u>**GAAAAAGTGGCACCGAGTCGGTGCTTTTTTT-3’, underlined bold sequences indicate the modified sequence and introduced class II terminator) through PCR mutagenesis with primers sgRNA-F (5’-ATCAGGCGCCATTCGCCATTCAGG-3’) and sgRNA-IITER-R (5’-AAAAAAAGCACCGACTCGGTGCCACTTTTTCAAACAGATAAC-3’) (Figure 4B). The IVT reaction (10 μl) containing 40 mM Tris-HCl pH 8.0, 16 mM MgCl_2_, 2 mM spermidine, 5 mM DTT, 0.2 μM inorganic pyrophosphatase, 1.5 U/μl RNase Inhibitor, 4 mM each of ATP, CTP, GTP and UTP, 150 nM T7 RNAP WT or S43Y mutant and 70 nM DNA template was carried out at 37°C for 1 h. After incubation, 1 μl reaction mixture was directly mixed with 2 μl 3x denaturing loading buffer and 3 μl ddH_2_O, heated at 80°C for 2 min, then the samples were loaded onto a 12% TBE PAGE gel and electrophoresis was conducted at 100 V for 1 hour. In addition to the artificial terminators, termination of WT T7 RNAP and S43Y mutant on several native class II terminators were also tested using DNA templates from plasmid pETsnrs-1 of which the T7 TΦ was replaced by designated native class II terminator sequence. The universal primers (pETsnrs-F and pETsnrs-1R) were used to amplify the templates, then IVT reaction was conducted as that for class I terminators above (Figure 4C).

The IVT of eGFP sgRNA and various sgRNA (Table S3) without terminator by T7 RNAP WT and S43Y mutant was also compared (Figure 5). DNA templates were amplified by PCR using a pair of universal primers sgRNA-F (5’-ATCAGGCGCCATTCGCCATTCAGG-3’) and sgRNA-R (5’-AAAAAAAGCACCGACTCGGTGCCACT-3’). IVT reaction condition was as mentioned above.

Overall IVT yield of WT T7 RNAP and S43Y mutant was compared on the production of tRNA^Ile^ (5’-GGCCGGTTAGCTCAGTTGGTTAGAGCGTGGTGCTAATAACGCCAAGGTCGCGGGTTCGATCCCCGTACTGGCCA-3’) and tRNA^Leu^ (5’-GGTAGCGTGGCCGAGCGGTCTAAGGCGCTGGATTAAGGCUCCAGTCTCTTCGGTTGCGTGGGTTCGAATCCCACCGCTGCCA-3’). IVT templates were constructed by replacing the sgRNA coding sequence in plasmid pUCT7sg-eGFP with the tRNA coding sequences. IVT reaction condition and analysis were as those for sgRNA above. The comparison of the IVT production of various long RNA (from 900 nt to 4100 nt) by WT T7 RNAP and S43Y was conducted in a 10 μl reaction containing 40mM Tris-HCl pH 8.0, 16 mM MgCl_2_, 2 mM spermidine, 5 mM DTT, 0.2 μM inorganic pyrophosphatase, 1.5 U/μl RNase Inhibitor, 4 mM each of ATP, CTP, GTP and UTP, 150 nM T7 RNAP WT or S43Y mutant and 20 ng/μl DNA templates. After incubation at 37°C for 1 h, 1 μl reaction mixture was mixed with 4 μl 3x denaturing loading buffer and 7 μl ddH_2_O, heated at 80°C for 2 min, each sample was then loaded onto a 1.5% TAE agarose gel and analyzed as mentioned above (Figure S4).

### RdRp assays

sgRNA products synthesized by T7 RNAP WT and S43Y mutant were purified with RNA purification kits (New England Biolabs) before analyzed by high performance liquid chromatography (HPLC) using PLRP-S column (4000 A, 8 μM, 4.6 × 150 mm) (Figure 5B). In order to demonstrate that the RdRp activity of T7 RNAP is reduced by the S43Y mutation, the run-off sgRNA produced by T7 RNAP S43Y IVT was further purified by collection the HPLC fraction of the major peak (Figure 5B) and used as the template for RdRp extension assay. The 10 μl reaction mixture contained 40 mM Tris-HCl pH 8.0, 16 mM MgCl_2_, 2 mM spermidine, 5 mM DTT, 0.2 μM inorganic pyrophosphatase, 1.5 U/μl RNase Inhibitor, 4 mM each of ATP, CTP, GTP and UTP, 150 nM T7 RNAP WT or S43Y mutant, and 500 ng/μl HPLC purified sgRNA. The RdRp extension reaction was incubated at 37°C for 1 h. Then 2 μl 3x denaturing loading buffer and 2 μl ddH_2_O were mixed with 2 μl reaction mixture and heated at 80°C for 2 min. Finally the mixture was loaded onto a 12% TBE PAGE and analyzed as mentioned above (Figure 5A).

### Termination efficiency calculation

The gray value of each band for transcripts was quantified with ImageJ software. For accurate quantification, each gray value was transferred into mole number based the molecular weight of corresponding RNA. Termination efficiencies were calculated as the molar ratio between terminated transcripts and the sum of terminated and run-off transcripts.

### Structural interpretation

The crystal structures of the T7 RNAP initiation complex (Protein Data Bank code 1QLN) and elongation complexes (Protein Data Bank codes 1MSW) were used to analyze the location of the mutation S43Y in a structural context. PyMOL software was used for visualization.

## RESULTS

### Evolution of T7 RNAP towards reduced termination efficiency at T7 TΦ

We developed a phage-assisted continuous evolution (PACE) system to generate T7 RNAP mutants (Figure 1A). PACE has been used for the evolution of diverse proteins toward desired property, such as polymerases and genome-editing proteins with improved activities and specificities (28,29). While PACE has not been used to evolve T7 RNAP to increase its read-through transcription, we consider that it would serve as an ideal method to explore the elements within single-subunit RNAPs that are involved in transcription termination.

**Figure 1.**
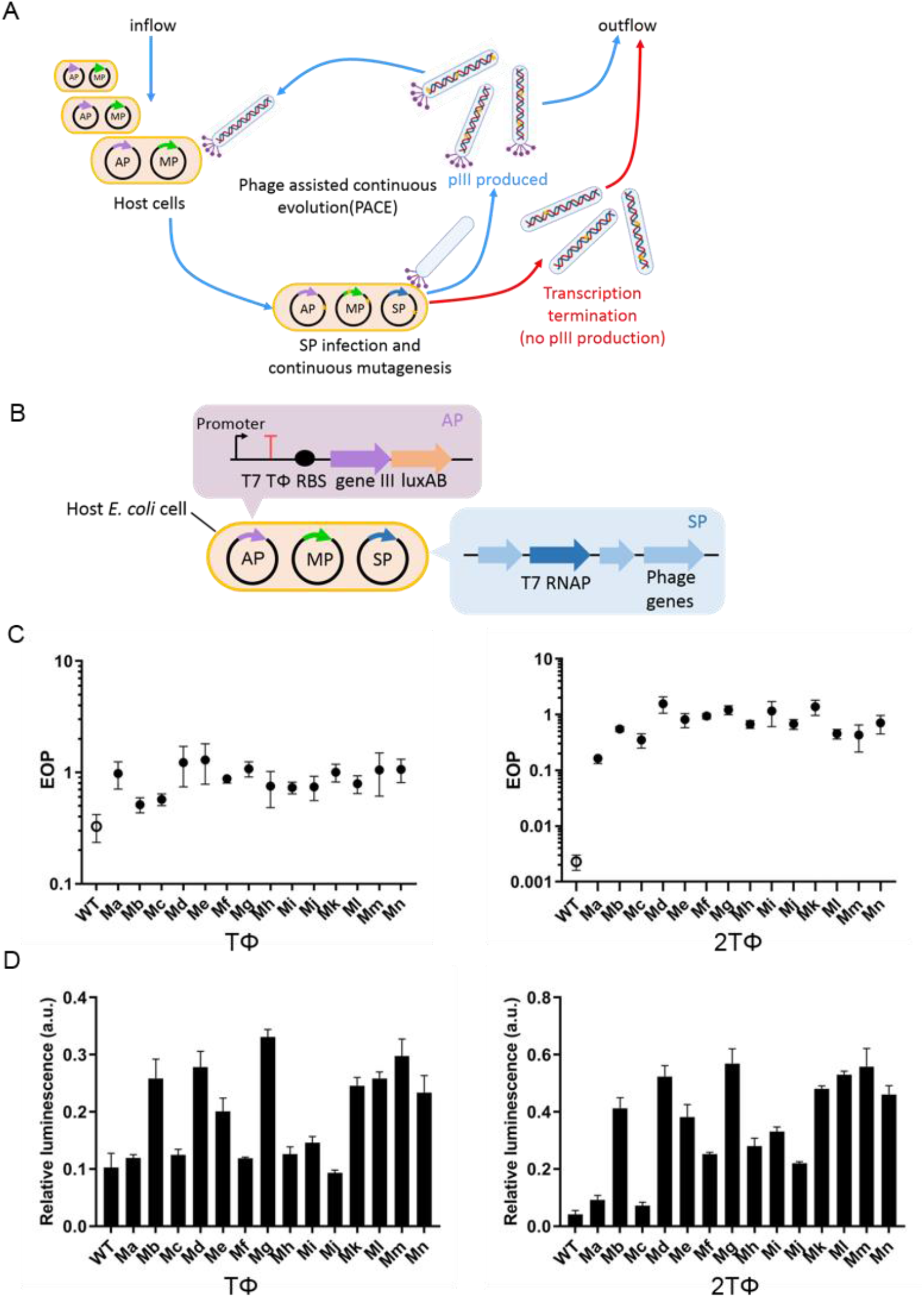
PACE of T7 RNAP for mutants with decreased termination efficiency at T7 terminator TΦ. **(A)** Schematic overview of the PACE in this study. The *E. coli* host cells contain two plasmids before infection: the accessory plasmid (AP), which correlates the termination efficiency of T7 RNAP to M13 phage propagation (details in (B)); and the mutagenesis plasmid (MP), which elevates the mutagenesis levels during PACE (27). The gene for pIII protein (gene III) in AP is under the control of a T7 promoter, while between the T7 promoter and gene III one or two tandem TΦ terminators were inserted to block the transcription of gene III by the T7 RNAP encoded in selection phage (SP) genome. Without pIII protein the M13 selection phage fails to infect host bacteria and gets washed out by the repeated dilution process. However, if the mutagenesis driven by the MP creates T7 RNAP mutants that can bypass the TΦ terminator, pIII protein is expressed and infectious progeny phages are produced to persist through the repeated dilution of PACE. **(B)** Key elements in AP and SP as described in (A). **(C)** Efficiency of plating (EOP) of selected phages indicated termination bypassing. EOP of the phages carrying T7 RNAP WT and selected mutants (Ma - Mn) was calculated by dividing the phage titer obtained on host cells carrying accessory plasmid pLATp1 (with one TΦ terminator inserted between T7 promoter and gene III, left panel) or pLATp2 (with two TΦ terminators inserted between T7 promoter and gene III, right panel) by that obtained on cells carrying pLATp3 (without TΦ between T7 promoter and gene III). **(D)** Termination bypassing of T7 RNAP WT and selected mutants (Ma - Mn) indicated by the transcription of luminescence reporter gene downstream of TΦ terminator. Relative luminescence was calculated by dividing the OD normalized luminescence obtained using host cells carrying reporter plasmid pLRTp1 (with one TΦ terminator inserted between T7 promoter and the reporter gene, left panel) or 2 (with two TΦ terminators inserted between T7 promoter and the reporter gene, right panel) by that obtained using cells carrying pLRTp3 (without TΦ between T7 promoter and the reporter gene).

In order to link the ability of T7 RNAP to bypass the terminator TΦ to the proliferation of phage M13, we replaced gene III in the phage genome with the T7 RNAP gene. While on an accessory plasmid pLATp1 carried by the host *E. coli* cells, TΦ was inserted between the T7 promoter and the ribosome binding site (RBS) in front of gene III (Figure 1B). We also constructed another accessory plasmid pLATp2 using two terminators of TΦ in tandem in order to exert stronger selection stress. It was reported that the production of infectious M13 phage particles increased with the increasing concentration of pIII (encoded by gene III) (30). Therefore, the ability of phages to propagate in the host cells carrying accessory plasmids is positively correlated with the level of gene III transcripts, which should only be produced by T7 RNAP bypassing the TΦ(s) upstream of gene III. Phages that persist through the repeated dilution process of PACE experiment are expected to produce evolved RNAP mutants with improved transcription activity in the presence of TΦ. As shown in Figure 1C, the efficiency of plating (EOP) of the evolved phage mutants significantly increased as compared with that of the WT, using either the accessory plasmid containing only one TΦ (1.5 to 4-fold increase) or that containing two (71 to 682-fold increase). This result suggested that the ability of the RNAPs encoded by these mutants to bypass terminators was improved. The effect of the evolution was also evaluated by another *in vivo* test of luminescence reporter assay. In the presence of only one TΦ, 8 out of the 14 mutants produced significantly stronger relative luminescence as compared with the WT, while in the presence of two TΦ in tandem, all the mutants produced relatively stronger luminescence (Figure 1D). Combining the results of the two *in vivo* tests, we inferred that mutations of key residues in the T7 RNAP involved in termination were generated, especially from the PACE when tandem terminators were applied.

### T7 RNAP mutants from PACE with reduced termination efficiency *in vitro*

We acquired the information of residues of 14 T7 RNAP mutants from PACE by sequencing (Figure 2A). We found that all mutants except Ma and Mc have a mutation at the 43^th^ residue S43, with residues of larger size including tyrosine (S43Y), phenylalanine (S43F) or leucine (S43L) replacing the original serine. We selected ten representative mutants (from Ma to Mj), overexpressed them in *E. coli*, and obtained purified T7 RNAP mutants. Alteration of these T7 RNAP mutants was directly tested in IVT with DNA templates harboring T7 promoter and terminator TΦ. The results showed that the termination efficiency on TΦ of all mutants except Ma and Mc was markedly decreased, and mutants (Md, Me, Mf and Mi) with most significant reduction in termination efficiency (over 20 %) all contain a common mutation S43Y (Figure 2B).

**Figure 2.**
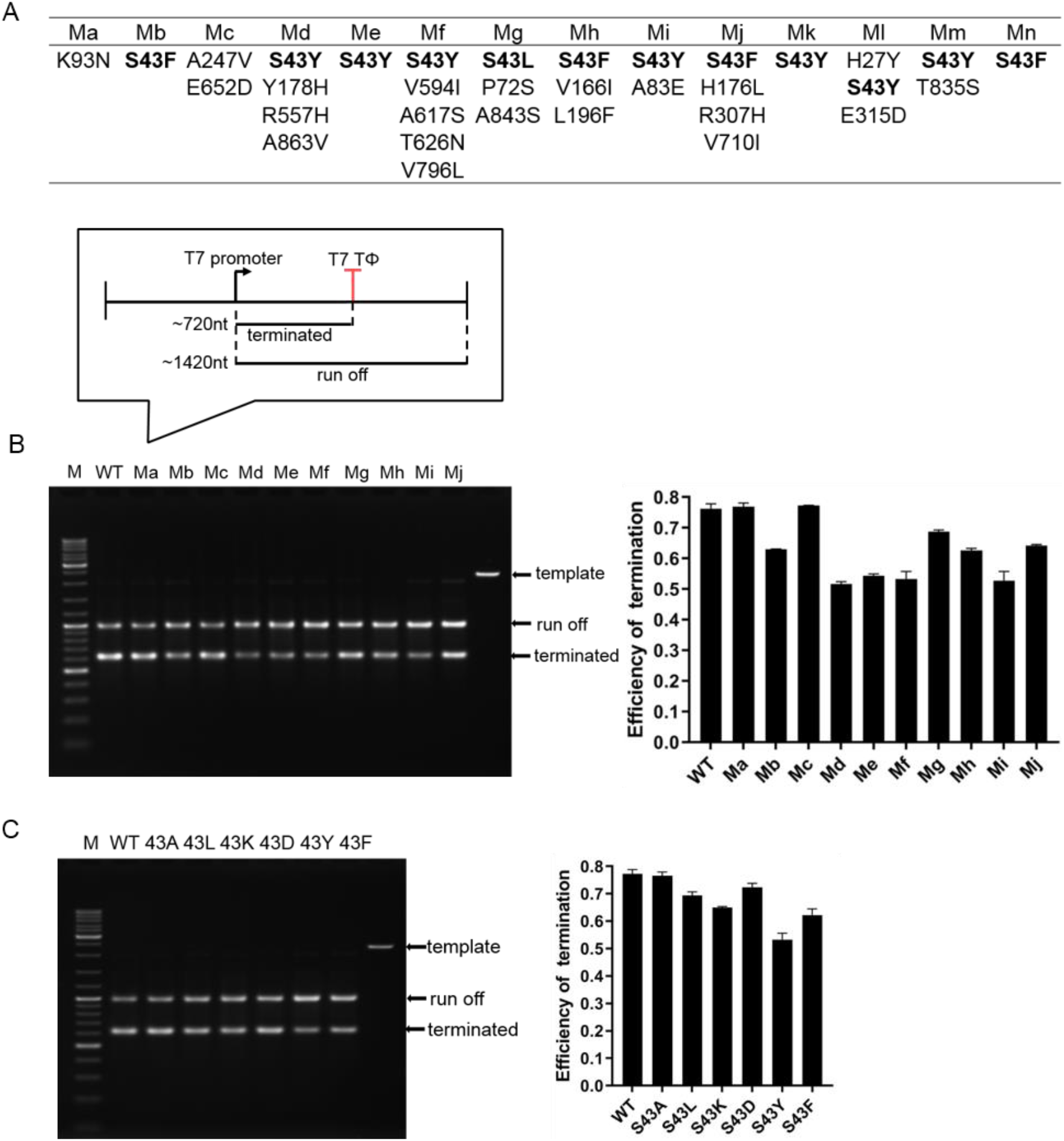
T7 RNAP mutants selected from PACE and their transcription termination *in vitro*. **(A)** Residues mutations of all potential termination-bypassing T7 RNAP mutants revealed by DNA sequencing. **(B)** Termination efficiency of WT and ten representative mutants (from Ma to Mj) of T7 RNAP on the DNA template containing T7 promoter and terminator TΦ (schematic showing the organization of DNA template and sizes of run-off and terminated transcripts on top of gel). Identity of DNA template and major RNA products was indicated by arrows. The termination efficiency of WT T7 RNAP or each mutant was calculated as the molar ratio between terminated transcripts and the sum of terminated and run-off transcripts from three independent assays and shown as the right panel. **(C)** Termination efficiency of WT T7 RNAP and mutants S43A, S43L, S43K, S43D, S43Y and S43F obtained with the same procedure as that for **(B)**.

We further investigated the effect of single mutation at the S43 position, constructing T7 RNAP mutants S43A, S43L, S43K, S43D, S43Y and S43F. Among these mutants, the S43Y mutant showed the greatest reduction on termination efficiency at T7 TΦ from nearly 78% to 53%. S43F, S43K and S43L reduced about 17%, 12% and 10%, respectively. The S43D mutant also has a slight reduction about 5% while the S43A mutation did not show any effect on termination (Figure 2C).

### S43Y mutation decreases the termination efficiency of T7 RNAP on class I terminators

We compared the termination efficiency of T7 RNAP WT and S43Y mutant in detail by varying the reaction conditions of IVT including enzyme concentration, template concentration and reaction time. The termination efficiency of S43Y had a slightly increase when the enzyme concentration was gradually increased, in contrast only fluctuations appeared for WT (Figure 3A). The termination efficiency of S43Y was slightly decreasing upon increasing template concentration, while the WT remained similar termination efficiency at various template concentrations (Figure 3B). Extension of reaction time has little effect on the termination of both the WT and S43Y mutant (Figure 3C).

**Figure 3.**
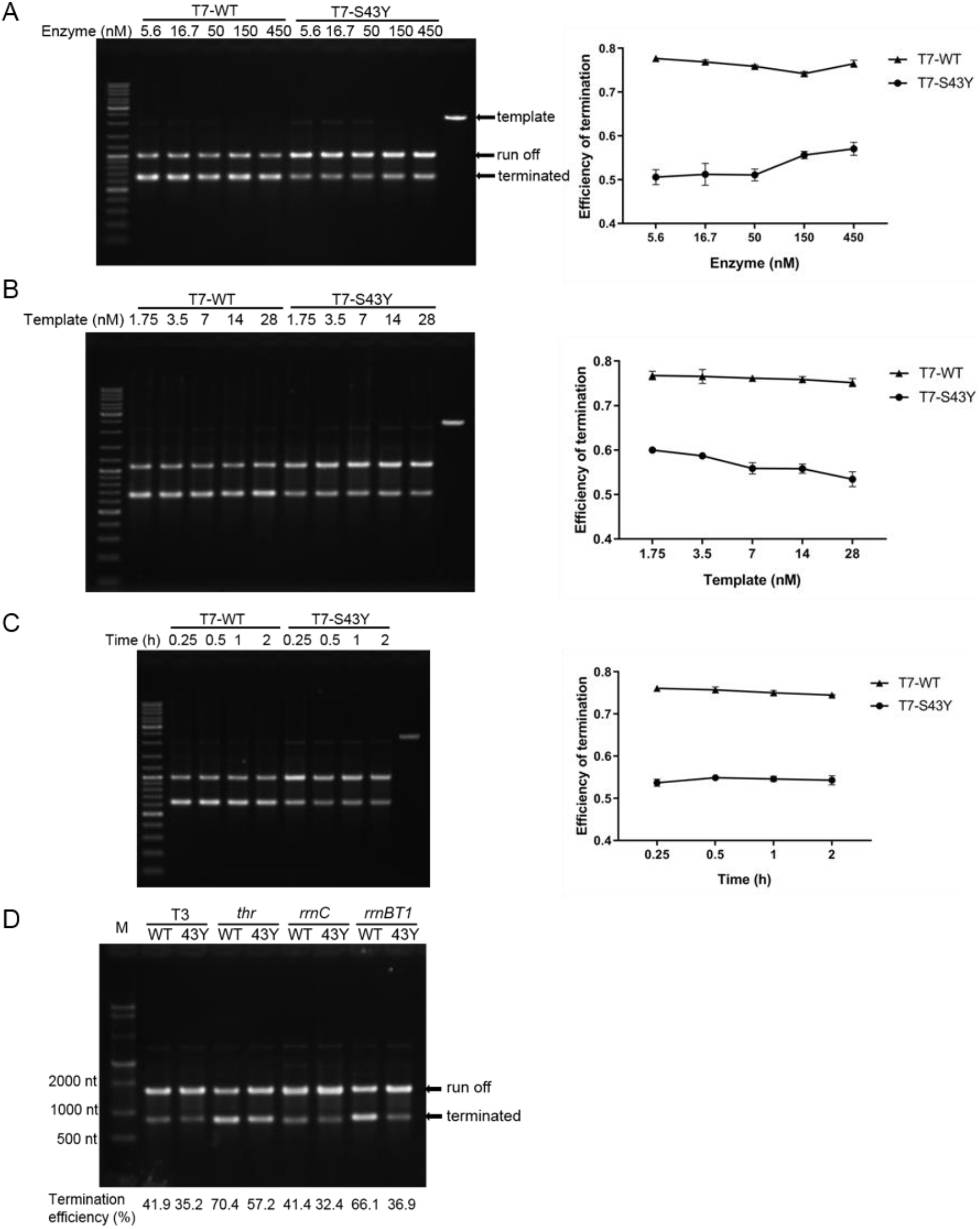
A detailed comparison between WT T7 RNAP and S43Y mutant on termination efficiency at class I terminators. **(A)** Termination efficiency of WT T7 RNAP and S43Y mutant at various enzyme concentrations (5.6 nM, 16.7 nM, 50 nM, 150 nM, and 450 nM). **(B)** Termination efficiency of WT T7 RNAP and S43Y mutant at various DNA template concentrations (1.75 nM, 3.5 nM, 7 nM, 14 nM, and 28 nM). **(C)** Termination efficiency of WT T7 RNAP and S43Y mutant at various reaction time (0.25 h, 0.5 h, 1 h, and 2 h). From **(A)** to **(C)**, the termination efficiency at each condition was quantified as the molar ratio between terminated transcripts and the sum of terminated and run-off transcripts from three independent assays and shown as graphs on the right. **(D)** Termination efficiency of WT T7 RNAP and S43Y mutant at four class I terminators: T3 TΦ, *thr* attenuator, *rrnBT1* terminator and *rrnC* terminator. The termination efficiency at each condition was shown at the bottom of corresponding gel lane.

The detailed comparison *in vitro* confirmed that S43Y mutation has a markedly effect on termination efficiency at T7 TΦ. T7 TΦ is a typical class I terminator for T7 RNAP. It has been reported that T7 RNAP also terminates at T3 TΦ and certain intrinsic terminators of bacterial RNAPs (11,31). To determine whether the S43Y mutation affects termination at other class I signals, we constructed IVT templates in which the T7 TΦ was replaced by T3 TΦ, *thr* attenuator, *rrnBT1* terminator and *rrnC* terminator, respectively. Compared to WT, the S43Y mutant showed obvious reduction on the termination efficiency at all terminators tested, suggesting a common effect on class I terminators (Figure 3D).

### S43Y mutation reduces the termination of T7 RNAP at class II terminators

As shown in Figure 3D, the reduction on termination efficiency of S43Y at the terminator rrnBT1 was the most significant compared to those at other class I terminators. It turned out that the rrnBT1 also harbors a class II terminator downstream of the class I terminator (8). Thus, although the S43Y mutation was obtained through evolution on class I structure-dependent terminator, it may also affect the termination at class II sequence-dependent terminators. We constructed templates with an artificial class II terminator (5’-ATCTGTTTTTTTTT-3’) inserted to compare WT T7 RNAP and S43Y mutant in IVT. The results showed that the termination efficiency of S43Y mutant was also decreased compared to WT at the class II signal (Figure 4A). The core sequence of a class II signal (5’-ATCTGTT-3’) inserted into the transcription template of an eGFP sgRNA (103 nt) resulted in the production of ~78 nt terminated IVT products by WT T7 RNAP (Figure 4B). However, the terminated products were eliminated by the S43Y mutation, meanwhile the run-off transcripts were increased significantly compared to that of WT (Figure 4B). Detailed comparison was also made in a range of reaction conditions including enzyme concentration, template concentration and reaction time, which further verified the effect of S43Y on the class II signal (Figure S2). Several natural class II terminators including signals in PTH, VSV, Adeno5 and the CJ site were also tested and the S43Y mutation consistently decreased the termination efficiency at all class II terminators tested (Figure 4C).

**Figure 4.**
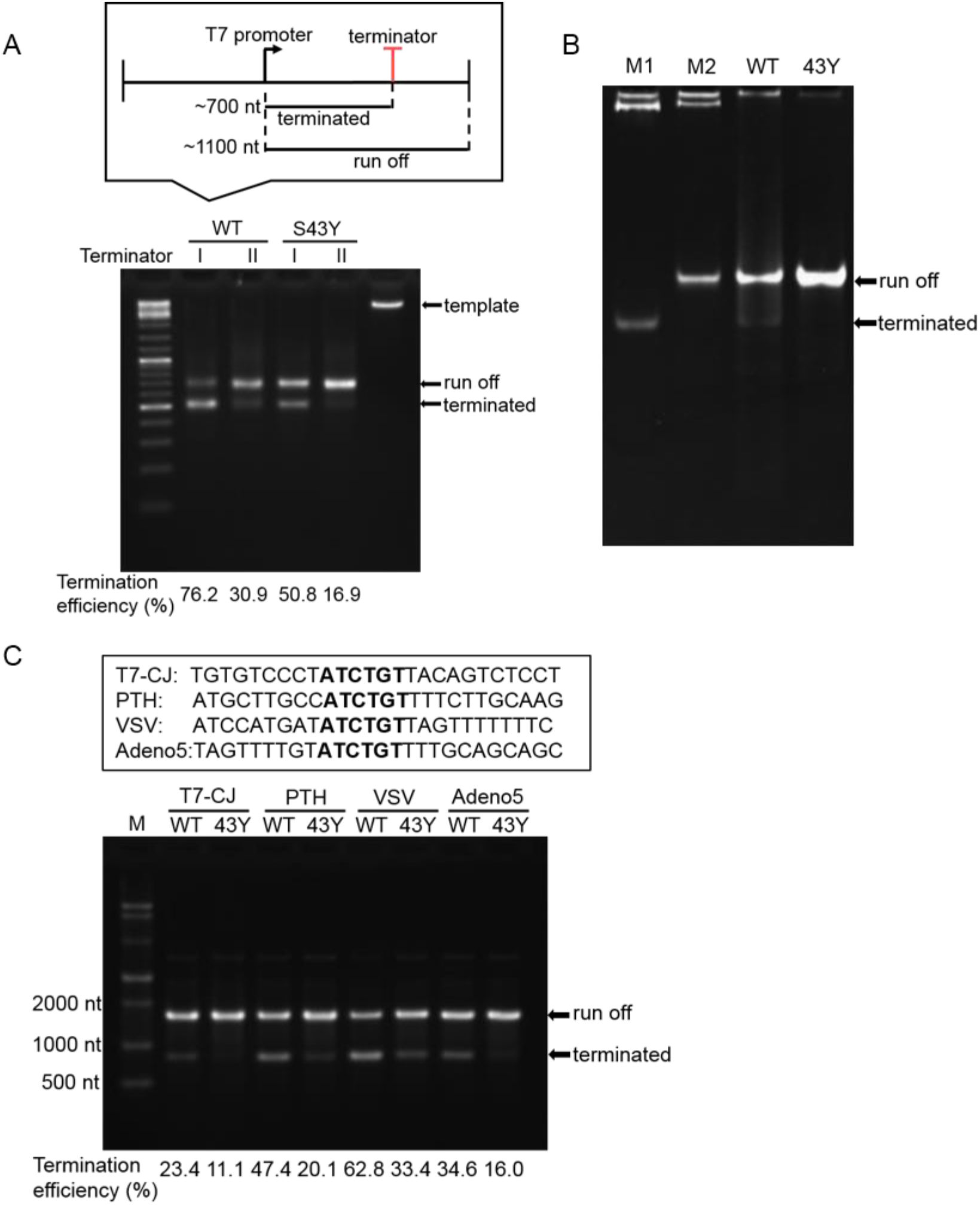
Termination of WT T7 RNAP and S43Y mutant at class II terminators. **(A)** Termination efficiency comparison between WT T7 RNAP and S43Y mutant on the templates containing either a class I (TΦ) or class II (5’- ATCTGTTTTTTTTT -3’) terminator (schematic showing the organization of DNA template and sizes of run-off and terminated transcripts on top of gel). Identity of DNA template and major RNA products was indicated by arrows. The termination efficiency at each condition was shown at the bottom of corresponding gel lane. **(B)** IVT of an eGFP sgRNA carrying an inserted class II termination signal (5’-ATCTGTTT-3’) by WT T7 RNAP and S43Y mutant. M1 is the 78 nt 5’ fragment of the sgRNA as a size marker for the expected terminated transcripts. M2 is 103 nt full-length sgRNA size marker. The terminated and run-off RNA gel bands were marked by arrows. **(C)** Comparison of the termination efficiency of WT T7 RNAP and S43Y mutant at various class II terminators (PTH, VSV, Adeno5 and the CJ site). The DNA sequences of the four natural class II terminators were shown on top of the gel, and calculated termination efficiency was shown at the bottom of each lane.

### S43Y mutation attenuates the RNA-dependent RNA polymerase activity of T7 RNAP

Beside the DNA-dependent RNA polymerase (DdRp) activity, T7 RNAP was found to retain the RNA-dependent RNA polymerase (RdRp) activity as well. T7 RNAP would catalyze RNA-primed RNA synthesis if the run-off RNA displays 3’-end complementarity to itself generating hairpin structure or to another RNA molecule forming intermolecular duplexes, yielding products longer than the run-off RNA (22,23,32,33). When we use WT T7 RNAP to synthesize the eGFP sgRNA (with terminal hairpin structure to mimic an RNA-primed RNA template structure), the 3’ overextended transcripts from the RdRp activity shown as gel bands and smears moving slower than the run-off product bands in PAGE gel were clearly observed (32,33). In this study when we investigate the termination efficiency of WT T7 RNAP and S43Y mutant on the class II terminator inserted in the eGFP sgRNA, we found that the S43Y mutation not only reduced the terminated products but also the overextended RNA products, resulting in significant increasing of run-off products (Figure 4B). This unexpected result led us to investigate the RdRp activity of the S43Y mutant in more detail. On IVT for the eGFP sgRNA without termination signal, the 3’ extended transcripts were obvious for WT T7 RNAP but barely observable from the S43Y mutant products (Figure 5A and Figure S3). HPLC analysis also showed additional peaks representing 3’ extended transcripts following the main peak corresponding to the run-off sgRNA from WT T7 RNAP products (Figure 5B). Consistent with the gel analysis, these additional peaks were not observed in the HPLC chromatogram for IVT products of S43Y mutant (Figure 5B). The DdRp and RdRp activities of T7 RNAP are mixed in IVT, thus we purified the run-off sgRNA and used it as the only template to compare the RdRp activity of WT T7 RNAP and S43Y mutant (33). Run-off sgRNA from T7 RNAP S43Y mutant IVT was purified by collecting the main peak (Figure 5B) after HPLC (Figure 5A lane 3) and served as an RNA-primed RNA template to test the 3’ extension from RdRp activity. Through PAGE gel analysis, we observed that the WT T7 RNAP extended the sgRNA judged from the emerging of additional slower-moving gel bands (Figure 5A lane 4), while much less of such bands were produced by the S43Y mutant (Figure 5A lane 5). Thus, the S43Y mutation attenuates both the transcription termination and RdRp activity of T7 RNAP, without disturbing its DdRp - the major activity for transcription.

**Figure 5.**
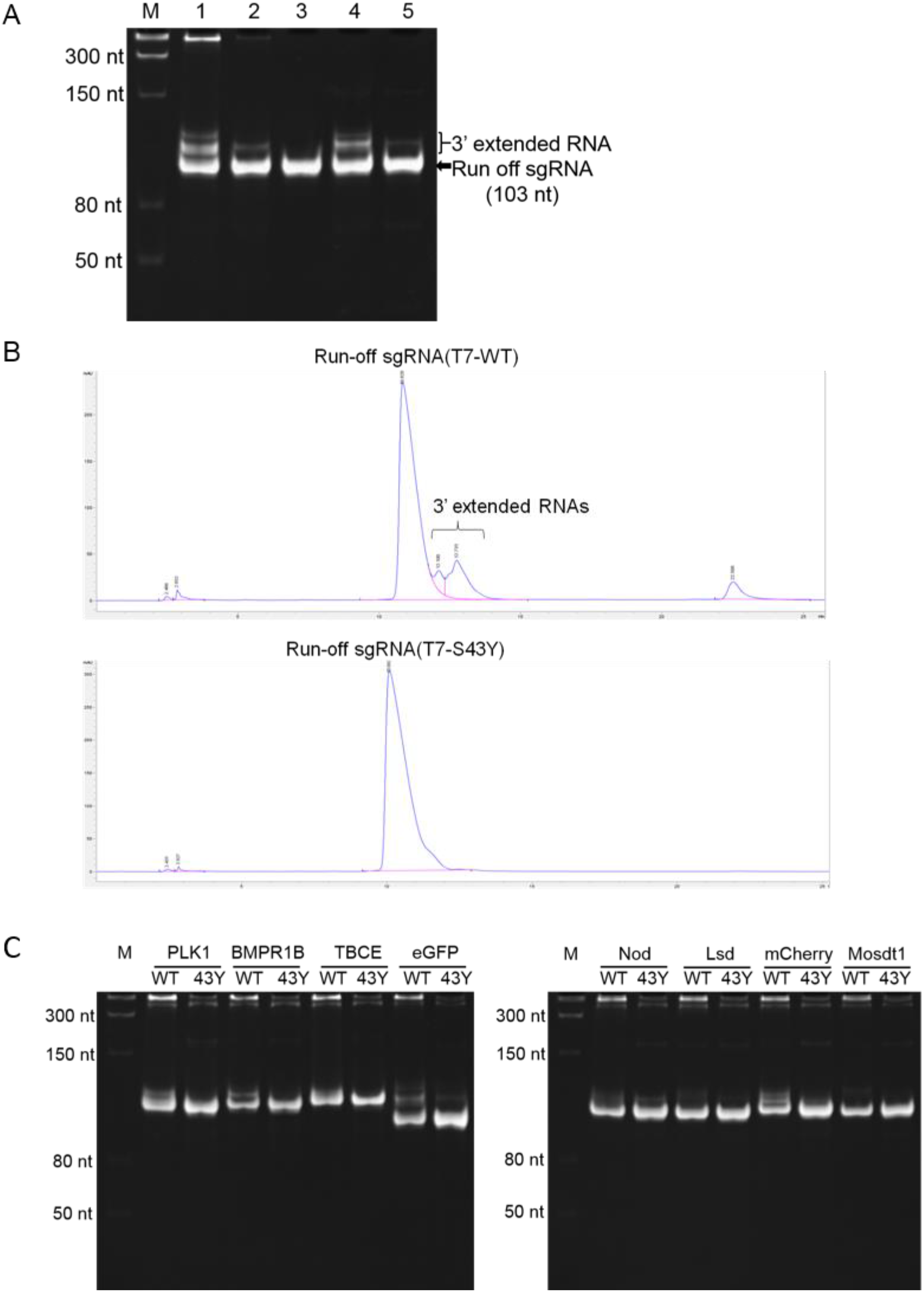
RdRp activity of WT T7 RNAP and S43Y mutant. **(A)** IVT products by WT T7 RNAP (lane 1) or S43Y mutant (lane 2) on DNA template encoding an eGFP sgRNA. The run-off sgRNA from S43Y products (lane 2) was purified by HPLC (lane 3). With the purified run-off sgRNA as reaction template, T7 RNAP WT produced significant amount of extended products moving slower than the precise sgRNA (lane 4), while such extension resulted from RdRp activity was much weaker for S43Y mutant as less extra products were produced (lane 5). For each lane a total 1 μg RNA was loaded. The 3’ extended RNA and run-off sgRNA were indicated at the right of the gel. **(B)** HPLC chromatogram of the eGFP sgRNA IVT products by WT T7 RNAP (top) and S43Y mutant (bottom). In the HPLC chromatogram additional peaks (extended products) following the main peak (run-off products) were obvious for WT T7 RNAP but not S43Y mutant. Corresponding RNA products were indicated on top of the chromatogram peaks. **(C)** IVT production of various sgRNA targeting different genes as indicated on the top by WT T7 RNAP and S43Y mutant.

We also compared the IVT production of multiple other sgRNAs targeting various genes by WT T7 RNAP and S43Y mutant. The 5’ 20 nt sequences of the sgRNA vary for targeting genes, and the rest 83 nt sequences of gRNA backbone are the same and are likely to form the same terminal secondary structure. Gel analysis demonstrated that for all the sgRNA tested, S43Y IVT provided significantly improved yield and purity compared to WT T7 RNAP IVT (Figure 5C). However, the IVT yield of RNA with other length and sequences by WT T7 RNAP and S43Y mutant is similar (Figure S4). Thus the high yield of sgRNA by S43Y mutant should be due to its reduction in RdRp activity but not enhanced DdRp efficiency.

## DISCUSSION

### S43 is a key residue of T7 RNAP involved in template interaction

It has been proposed that the transcription termination of T7 RNAP involves a reverse process of the transition from transcription initiation to transcription elongation (9). When T7 RNAP undergoes transition from the initiation complex (IC) to the elongation complex (EC), its N-terminal domain (residues 1 to 325) which is involved in promoter recognizing and DNA melting substantially alters conformation while the other portion of the enzyme has little change, as revealed in the complex structures (7,34). During the transition, a significant change is the rotation of the C2-helix (residues 46-55) and its stacking on the C1-helix (residues 28-41) to form an elongated C-helix (Figure 6A). The C-helix (Residues 28-71) replaces the promoter binding domain (PBD) in the IC and aligns with the template DNA strand in the EC (Figure 6A). Therefore, the refold of C-helix cooperated with the rotation of PBD reconstructs the T7 RNAP and facilitates the formation of EC (35). It was also suggested that the formation of C-helix is important for the stability of EC, as disrupting the refold of C-helix through mutagenesis of residues within this helix to proline increases abortive transcripts and prevents stable EC (36). As shown in Figure 6A, the side chain of Arg^50^ in C2-helix is only 4.4 Å away from the template-strand DNA in EC structure. And a positive-to-negative charge mutation R50E resulted in the accumulation of abortive transcripts during T7 RNAP transcription (36), indicating that R50 may directly interact with the DNA template and the C2-helix plays a crucial role in stabilizing the association between enzyme and DNA. Ser^43^, on the other hand, not noticed previously and to our knowledge the only residue reported in T7 RNAP on which a single mutation specifically reduces termination, locates in the loop between the C1-helix and C2-helix like a hinge, suggesting its potential role to regulate the formation of the C-helix and the alignment between the C2-helix and template DNA strand. By this means S43 regulates the enzyme-DNA interaction indirectly. As we observed, mutations of Ser^43^ to Leu, Lys, Asp, Tyr and Phe except for Ala all attenuated the termination of T7 RNAP (Figure 2C), presumably because that these larger residues at the hinge position alter the flexibility of the C2-helix, resulting in tighter DNA template association to reduce the disassociation of enzyme from DNA during termination. The Tyr and Phe mutations of Ser^43^ exhibited the most significant effect, possibly due to a direct involvement of their large side chain and/or the aromatic ring to interact with the DNA. Meanwhile, when interacting with RNA template in the RdRp mode of T7 RNAP, the S43 should play similar role in control the strength of template binding. Although the S43Y mutation enhances DNA association of T7 RNAP, the subtle conformation change of the C-helix that favors DNA binding may lead to clash with RNA template to abolish the RdRp activity as we have shown.

**Figure 6.**
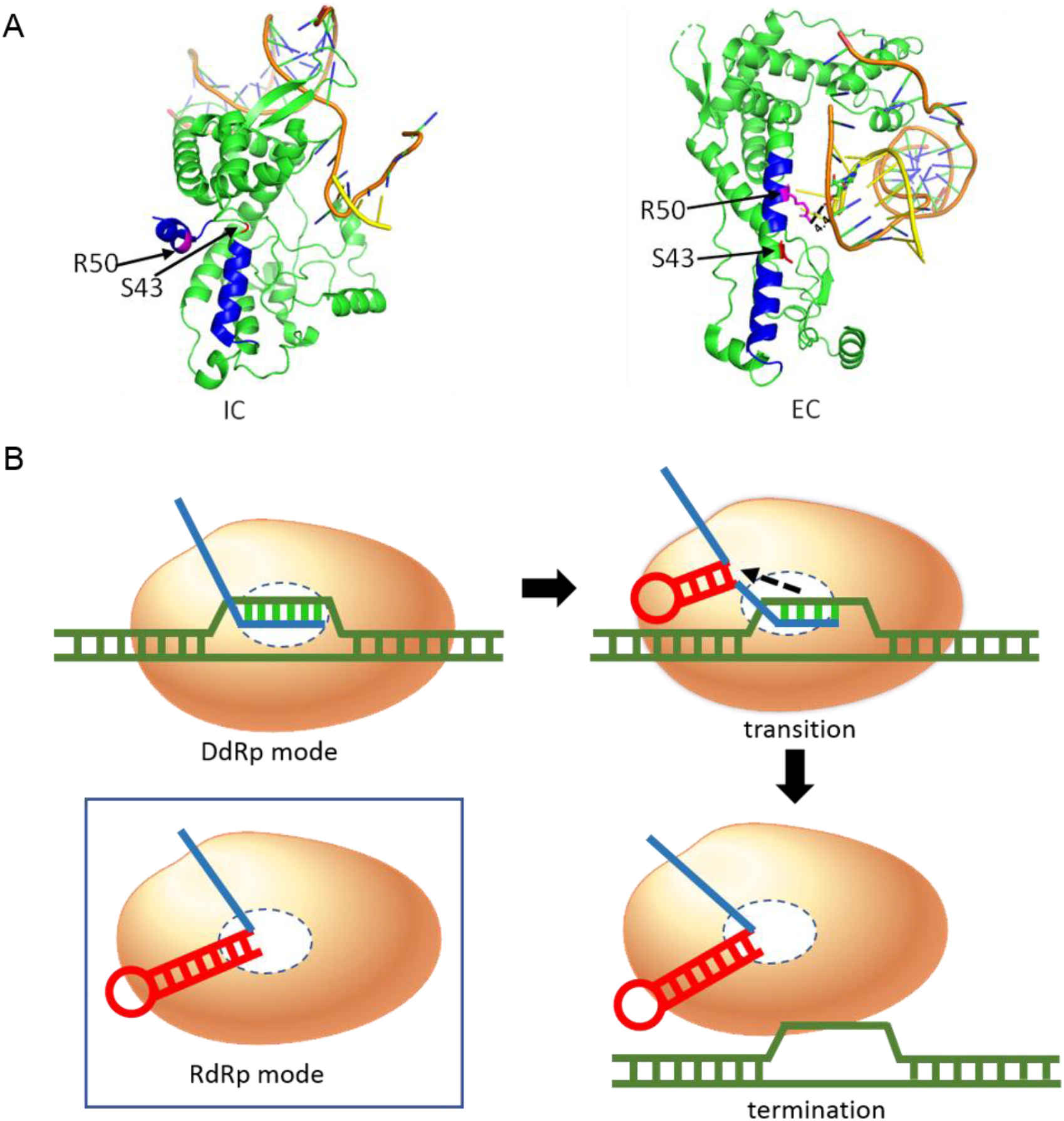
Role of S43 in the initiation complex (IC) and elongation complex (EC) structures of T7 RNAP and the proposed mechanism of termination at class I terminator. **(A)** The N-terminal domain (residues 1 to 325) of T7 RNAP in the IC (left, PDB accession no. 1QLN) and EC (right, PDB accession no. 1MSW) structures, as well as the DNA template and RNA transcript were shown. S43 (colored in red) as a hinge of C1 helix and C2 helix (in blue) and R50 (in purple) in C2 helix were indicated by arrows. The shortest distance between R50 and the DNA template was measured and shown. **(B)** Transition from DdRp into RdRp mode causes transcription termination of T7 RNAP at class I stem-loop terminator. During the transcription elongation, T7 RNAP (in orange) exhibits DdRp activity (active site shown as the dotted circle) to produce RNA transcript (in blue) on DNA template (in green). At a class I terminator, the formation of a stable stem-loop structure in nascent RNA may impede the stability of the DNA/RNA dulex in the active site and lead to competition between the DdRp and RdRp activity of T7 RNAP. If the fully formed stem-loop RNA structure triggers the RdRp activity of T7 RNAP as shown in the box, the enzyme disassociates from the DNA template and transcription is terminated.

### Mechanism of the transcription termination of T7 RNAP at class I terminators

As shown in Figure 6B, the DdRp mode presents the transcription elongation complex of T7 RNAP, in which the nascent RNA forms a short duplex with DNA template and is extended in the active site. In the process of transition from transcription elongation to termination, the nascent RNA would generate a hairpin structure at the class I terminator which causes EC collapse with RNA and polymerase released from the DNA template. The class I termination of T7 RNAP was considered to have a common mechanism with the termination at the intrinsic termination signals by bacterial RNAPs and several models are proposed concerning the mechanism of such kind of termination triggered by hairpin terminator structure in nascent RNA. Since it was revealed that the 5’ end of the emerging nascent RNA would interact with the N-terminal domain of T7 RNAP, and this RNAP-RNA interaction was proposed to regulate the stability of EC (37,38), these models could be classified into three categories while commonly focused on the steric clash of the RNA stem-loop with the exit pore of RNA transcripts in the polymerase (39). Most models implied the alteration of the active site and the disruption of RNA: DNA hybrid in upstream region (39,40). The translocation of RNA polymerase was also suggested to be essential for RNA release at the intrinsic terminator (41). The termination was reported to occur in two steps including RNAP pausing and the following RNA release (42). And the U-rich tract with low thermodynamic stability followed the hairpin-structure was suggested to contribute to the pausing and termination (40,43). Such pausing was supported by the effect of enzyme and template concentrations on the termination efficiency of S43Y (Figure 3A and Figure 3B) in our work. The results also indicated that the collision of multiple RNAP molecules at the termination site is more significant for S43Y mutant, and further indicated a longer pausing of the S43Y mutant at termination site compared to that of the WT T7 RNAP.

In this study, the S43Y mutation attenuates the class I termination and the RdRp activity simultaneously, giving us insight into the connection between transcription termination and the RdRp activity of T7 RNAP. And considering the formation of a stem-loop RNA structure as the common and key feature in both situations (Figure 6B), we proposed a model in which the formation of a stem-loop RNA structure at the class I terminator triggers the RdRp acitivity of T7 RNAP, and the competition between the RdRp and DdRp, in addition to previously suggested steric clash between the RNA stem-loop and the exit pore of polymerase, causes the pausing of the RNAP. Consequence of such competition depends on the context of sequences at and downstream of the terminator, which determines the stability of the RNA stem-loop in the terminator, and the stability of the RNA/DNA duplex in the active site of the RNAP. Providing a U-rich tract downstream of the terminator, the stability of the RNA/DNA duplex in the active site of RNAP should be weaken, thus favoring the competition towards the RdRp mode and transcription termination. On the other hand, alterations in the RNAP that enhance DNA template binding but weaken RNA template binding, such as the S43Y mutation, would favor the competition towards the DdRp mode and continuous transcription elongation.

The class II terminators share common sequence (5’-ATCTGTT-3’) in non-template strand DNA, such signals do not form obvious hairpin structure and the incorporation of IMP destabilizing the secondary structure of RNA was reported no effect on such termination, indicating a sequence-specific termination manner (44). The mechanisms underlying the two classes of termination were believed to be different. As a modified and less processive form of T7 RNAP which was proteolytically cleaved between residues 178 and 179, and several other mutants of T7 RNAP with alterations in this region such as R173C, failed to recognize class II terminators but remained robust termination at class I terminators (45,46). However, common features were also existing for the two types of termination. For example, although the class II termination is sequence-specific and orientation-dependent, the U-rich tract downstream also facilitates termination (8,10), and in our study, the two types of termination were simultaneously reduced by the mutation S43Y, suggesting that the stability of the RNA/DNA duplex in the active site of RNAP and the binding strength of the RNAP to DNA template affect the termination at both classes of terminators.

### A novel enzymatic reagent for IVT

With both transcription termination and RdRp activity of T7 RNAP attenuated by the S43Y mutation, the two major undesired byproducts from IVT - terminated transcripts and 3’ overextended transcripts are significantly reduced (Figures 3–5). Meanwhile, the S43Y mutant maintains the high yield of WT T7 RNAP (Figure S4). Introduction of a single mutation into the widely used T7 RNAP without changing the IVT formula provides a simple solution to improve the purity of IVT products.

## SUPPLEMENTARY DATA

Supplementary Data are available online.

## ACKNOWLEDGEMENT

We thank all lab members for helpful discussion.

